# ORBIT: Annotation-Aware Empirical Enrichment and Semantic Reranking for Interpretable Functional-Class Recovery

**DOI:** 10.64898/2026.07.01.735870

**Authors:** Benjamin L. Kidder

## Abstract

Gene-set interpretation workflows are widely used to summarize transcriptomic and proteomic experiments, yet standard enrichment tools often return long, redundant result tables that require substantial manual consolidation. We developed ORBIT (Ontology-Ranked Biological Interpretation Tool), an annotation-aware interpretation workflow that combines empirical enrichment, semantic reranking, and redundancy-aware representative-term selection to prioritize interpretable functional summaries from gene sets.

We evaluated ORBIT on a curated tiered benchmark of human functional-class gene sets spanning clean reference sets, size-ladder variants, and mixed-difficulty cases. On the 45-set core benchmark, ORBIT semantic achieved higher expected-class recovery than Enrichr and PANTHER Gene Ontology molecular-function baselines, with a mean reciprocal rank of 0.916 and top-1 recovery of 0.889. Bootstrap confidence intervals and paired permutation testing supported the robustness of this advantage, and supplemental analyses extended the comparison to g:Profiler. In a GPCR mixed-function case study, ORBIT compressed redundant enriched terms into semantic representative neighborhoods, illustrating how long enrichment outputs can be converted into reviewable biological summaries.

We applied ORBIT across complementary transcriptomic analyses spanning single-cell immune-cell markers, interferon-beta stimulation, and breast-cancer subtype biology. In these settings, ORBIT condensed marker and differential-expression signatures into prioritized functional summaries with explicit links to the genes, cell types, and annotation classes supporting each result. Together, these applications show how ORBIT can move from ranked enrichment terms to structured biological interpretation, preserving the evidence trail from input genes to functional summaries.

## INTRODUCTION

Functional interpretation of gene lists remains one of the central tasks in modern genomics and proteomics. As large-scale experiments routinely produce hundreds to thousands of candidate genes, investigators depend on enrichment analysis and ontology-driven annotation resources to translate those lists into biological hypotheses. The Gene Ontology (GO) was originally created to provide a structured, computable vocabulary for molecular function, biological process, and cellular component, enabling cross-dataset and cross-species functional reasoning at scale^1–3^. That resource has since expanded in both scope and annotation depth, becoming a foundational substrate for enrichment and functional interpretation workflows.

This problem has been recognized since early GO-based interpretation tools such as Onto-Express, GOstat, DAVID, and related ontological analysis frameworks began to scale secondary interpretation beyond manual inspection^4–8^. Widely used modern interfaces such as Enrichr, WebGestalt, clusterProfiler, and STRING provide broader library coverage and more convenient ranked output, but still leave users to consolidate overlapping terms and decide which ranked result best captures the dominant biological theme^9–15^. General enrichment-analysis best practices emphasize that term redundancy, library choice, and post hoc interpretation are major determinants of how useful an enrichment result will be to end users^16–22^.

These challenges are especially relevant when the goal is not simply to obtain any enriched term, but to recover an interpretable, class-level biological explanation for a gene set. In this context, semantic relationships among ontology-linked terms matter alongside enrichment statistics. The broader gene-set analysis literature has repeatedly shown that enrichment results depend strongly on statistical formulation, gene-set definition, and the handling of correlated or overlapping categories^16–22^. Semantic similarity analyses in GO have long been recognized as useful for comparing functional relatedness, prioritizing coherent interpretations, and reducing ontology-driven redundancy ^23–25^. However, many commonly used enrichment workflows do not explicitly integrate semantic coherence and query-level consensus as secondary ranking signals layered on top of an empirical enrichment backbone.

ORBIT (Ontology-Ranked Biological Interpretation Tool) was developed to address this gap by adding an annotation-aware prioritization layer to gene-set interpretation. ORBIT combines empirical enrichment, annotation-derived feature construction, semantic reranking, and redundancy-aware clustering to prioritize functionally coherent terms and representative class-level summaries. The workflow draws on curated offline resources that are central to current functional interpretation practice, including UniProt, GOA, QuickGO, Ensembl, and the Human Protein Atlas^26–31^. The motivation for the method is practical as much as statistical: users often rely on the first few reported terms, and therefore a tool that elevates the most interpretable and biologically coherent class-level explanation can be more useful than one that returns only a long ranked list.

In this study, we benchmarked ORBIT against functional-enrichment baselines from Enrichr, PANTHER, and g:Profiler using a curated tiered benchmark of human functional-class gene sets. We then used a significant GPCR mixed-hard case study to illustrate how ORBIT organizes, prioritizes, and summarizes related terms into a more interpretable functional landscape, and extended that interpretability analysis with additional curated transcription-factor, kinase, and secreted-factor examples. Finally, we applied ORBIT to single-cell PBMC marker genes, IFNB-stimulated PBMC differential-expression signatures, and TCGA-BRCA tumor-subtype signatures to evaluate its behavior across common transcriptomic interpretation settings. Our goal was to test whether an annotation-aware empirical-plus-semantic workflow improves first-rank recovery and practical interpretability relative to standard external baselines on a benchmark tailored to simplified functional-class recovery, while also producing reviewable functional summaries in applied gene-expression use cases.

## MATERIALS AND METHODS

### Overview

The benchmarking framework used here compared ORBIT against primary molecular-function-oriented external baselines from Enrichr and PANTHER, with additional comparator analyses using g:Profiler, on a tiered benchmark of curated human functional-class gene set ^9–11^.

### Gene and Annotation Background

The ORBIT workflow operates on an offline human background of mapped proteins and associated annotation evidence. Gene-level representations integrate simplified function labels, localization context, ontology-linked evidence, confidence-weighted annotation text, and related background-protein context^26–31^. Term-level representations combine term labels, lexical and ontology context, co-annotation structure, and local evidence from the background annotation graph^23–25, 32^. These derived representations are used only for secondary prioritization and clustering; they do not replace the primary empirical enrichment statistics.

### ORBIT Enrichment Workflow

For a given input gene set, ORBIT first maps the query genes to the offline species background. It then evaluates enrichment against candidate terms using breadth-matched empirical null sampling over the same background. In the benchmarked configuration, primary significance estimates were derived from empirical permutation testing, followed by false-discovery-rate correction. This places ORBIT in the broader family of gene-set and pathway enrichment methods that extend early over-representation and gene-set scoring frameworks such as GOstat, GSEA, PAGE, and related model-based approaches^5, 17, 18, 33^. The main manuscript runs used semantic reranking and semantic clustering in addition to the primary empirical enrichment backbone.

The semantic reranking stage combines token-level coherence, embedding-level coherence, gene-term alignment, and query-level functional consensus. Token-level coherence is calculated from TF-IDF representations of query annotation text and term documents. Embedding-level coherence uses a deterministic 96-dimensional hashed dense projection of the same TF-IDF token space rather than an external large language model. Gene-term alignment averages cosine similarity between the proteins supporting the query and the candidate term document. Query-level functional consensus compares candidate term labels with the confidence-weighted distribution of simplified function classes in the mapped query genes. These semantic features are used only for secondary prioritization and redundancy reduction; they do not replace the empirical p-value and FDR backbone.

The semantic reranking score is computed as a multiplicative prioritization score. ORBIT first defines feature-specific multipliers and then combines them with the empirical significance score and source prior:

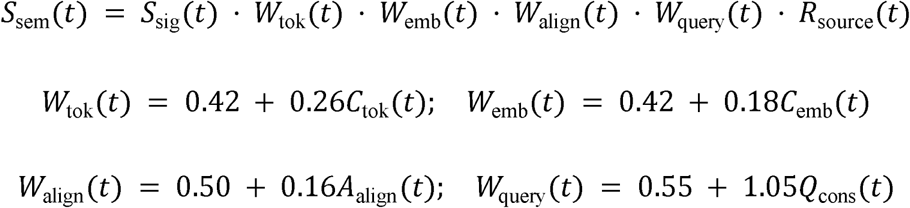

Here, C_tok and C_emb denote token and embedding coherence, A_align denotes gene-term alignment, Q_cons denotes query functional consensus, and R_source denotes the source-prior multiplier.

The source prior used in the manuscript configuration was 1.55 for ORBIT simplified-function labels, 1.18 for Gene Ontology molecular function, 0.96 for Gene Ontology biological process, and 0.90 for Gene Ontology cellular component. Redundant enriched terms are then assigned to semantic neighborhoods using a greedy representative-first clustering procedure with a default pairwise similarity threshold of 0.58. Pairwise similarity combines supporting-gene Jaccard overlap, sparse TF-IDF cosine similarity, dense hashed-vector cosine similarity, lexical overlap, prefix agreement, and source agreement. The first high-confidence ranked term initiating a cluster is retained as the representative term.

For a candidate term t, ORBIT estimates an empirical p-value from breadth-matched null permutations as:

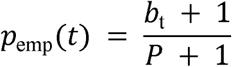

where b_t is the number of null permutations with at least the observed number of query hits and P is the number of permutations. Expected counts are averaged across permutations, and fold enrichment is calculated with a 0.5 pseudocount:

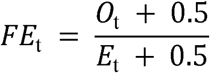

The empirical composite score used by the enrichment backbone is:

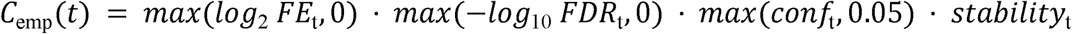

In the command-line implementation, the default enrichment setting uses 1,000 breadth-matched permutations, term-size bounds of 3 to 2,000 background proteins, and a fixed random seed for reproducibility.

### Benchmark Dataset Construction

Benchmarking was performed on a curated set of human functional-class gene sets organized into tiered benchmark bundles. The benchmark included clean reference sets, size-ladder variants, and mixed-difficulty sets. The core tier contained 45 non-secondary benchmark sets, while the exploratory tier extended this set to 52 entries by including additional secondary-reference cases.

Each benchmark gene set was associated with an expected simplified functional class, such as GPCR, receptor, kinase, phosphatase, transporter, transcription factor, RNA-binding protein, secreted factor, adhesion molecule, scaffold/adapter protein, chromatin regulator, or enzyme. Benchmark performance was evaluated by determining how highly each method ranked a term matching the expected class, using benchmark alias matching rather than exact string equality alone.

### External Baseline Workflows

#### Enrichr

Enrichr was benchmarked using a batch R workflow based on the enrichR package. The main Enrichr libraries evaluated were chosen to reflect current functional-library availability and practical relevance for the curated benchmark task. In practice, the most informative Enrichr external baseline was GO_Molecular_Function_2025. Other tested Enrichr libraries, including biological-process, cellular-component, and older Panther-linked libraries, were retained for context but were not the primary manuscript baseline because their recovery of the expected simplified functional classes was substantially weaker^12–15^.

#### PANTHER

Direct PANTHER benchmarking was performed using an R workflow built around rbioapi, which queried PANTHER functional datasets programmatically and exported result tables for downstream comparison. The strongest direct PANTHER baseline was PANTHER_GO_Molecular_Function, with additional PANTHER datasets retained for context and supporting analyses^10, 11^.

#### g:Profiler

A supplemental external comparator analysis was performed with g:Profiler/g:GOSt using the public g:Profiler API for Homo sapiens queries. Each benchmark gene set was submitted in two modes: a Gene Ontology molecular-function-only mode and an all-source g:GOSt mode. Raw JSON responses were archived, normalized into term tables, and passed through the same expected-label rank-recovery pipeline used for Enrichr and PANTHER. The g:Profiler analysis was included as a supplementary comparator to broaden external benchmarking beyond the two primary manuscript baselines while preserving Enrichr and PANTHER as the main molecular-function-oriented comparisons^34^.

#### External Result Harmonization and Comparison

External enrichment outputs were converted into manifest files and processed through a common comparison pipeline. For each benchmark gene set and external library, the comparison workflow parsed the ranked output terms, matched them against expected benchmark aliases, and recorded the best recovered rank. This created a unified benchmarking framework in which ORBIT, Enrichr, PANTHER, and the supplemental g:Profiler outputs could be compared on the same expected-label task. Pathway and ontology resources used in the broader comparison context included GO, KEGG, Reactome, STRING, DAVID, WebGestalt, clusterProfiler, and g:Profiler; not all resources were retained as primary manuscript comparators because the primary claim was intentionally scoped to representative molecular-function baselines and audited supplemental comparators^1–3, 14–17, 25, 32, 34, 35^.

#### Performance Metrics

Benchmark performance was summarized using mean reciprocal rank (MRR), top-1 recovery rate, top-5 recovery rate, top-10 recovery rate, and median recovered rank. These metrics were calculated independently for the core and exploratory benchmark scopes and, when useful, stratified by benchmark tier. We emphasized first-rank and top-k recovery because the practical purpose of ORBIT is to surface an interpretable class-level explanation within the first few reported terms rather than to optimize only the presence of a broadly related term somewhere in a long list.

Uncertainty estimates were added for the revised analysis by bootstrap resampling benchmark gene sets within each benchmark scope. For each method and metric, 10,000 bootstrap replicates were used to estimate 95% confidence intervals. Paired comparisons between ORBIT semantic and the strongest external Enrichr and PANTHER molecular-function baselines were performed using a paired sign-flip permutation test on per-gene-set metric differences, with 50,000 permutations. MRR was treated as the primary paired-effect metric because it preserves rank information and is less discretized than top-1, top-5, or top-10 recovery, which are binary at the individual gene-set level. Existing benchmark-tier labels were used for stratified MRR summaries; no post hoc biology categories were invented for this purpose.

#### Main Interpretability Case Study

To complement the aggregate benchmark analysis, a larger significant case study was selected from the tiered benchmark collection. The chosen example was the human GPCR mixed-hard query, which contained 12 input genes, produced 168 tested terms, and yielded 52 FDR-significant terms in the manuscript configuration. This case was used to generate representative-term plots, semantic-landscape figures, semantic-cluster size summaries, and representative-term overlap heatmaps.

The overlap heatmap summarizes Jaccard overlap among the supporting query-gene sets of cluster-representative enriched terms. It is an overlap-of-support plot rather than a semantic-similarity plot. The semantic-landscape figure shows significant enriched terms positioned by semantic coherence and statistical significance, with representative terms labeled for interpretability.

#### Additional Curated Example Analyses

To test whether the interpretability behavior generalized beyond receptor-heavy biology, we also evaluated four curated examples aligned to the benchmark class vocabulary: GPCR, transcription factor, kinase, and secreted factor. Each example was run through the same ORBIT enrichment workflow and summarized using semantic cluster counts in addition to raw significant-term counts, because many enriched terms represent partially redundant ontology variants.

The validated example outcomes in the current workspace were as follows: the GPCR example yielded 43 significant terms and 21 semantic clusters with GPCR as the top semantic term; the transcription-factor example yielded 19 significant terms and 13 semantic clusters with transcription factor as the top semantic term; the kinase example yielded 45 significant terms and 17 semantic clusters with kinase as the top semantic term; and the secreted-factor example yielded 71 significant terms and 25 semantic clusters with secreted factor as the top semantic term. We additionally generated a dedicated transcription-factor case-study output folder and figure bundle using the same figure-generation workflow used for the GPCR example.

#### Seurat PBMC3K single-cell demonstration

To demonstrate use downstream of single-cell RNA-seq clustering, we analyzed the SeuratData PBMC3K example with standard Seurat-style cell-type annotations and marker sets. Marker genes from nine annotated PBMC populations were passed to the ORBIT downstream annotation workflow to generate marker-to-biology summary tables, localization summaries, and cell-type-by-function enrichment dot plots. This demonstration was designed as a biological face-validity analysis: expected cell-type functions were prespecified from canonical PBMC biology and then compared with ORBIT-enriched marker annotations.

#### IFNB-stimulated PBMC expression use case

To evaluate ORBIT in a perturbational expression-analysis setting, we analyzed the SeuratData IFNB-stimulated PBMC dataset. Differential expression was computed for IFNB-stimulated versus control PBMCs overall and within annotated cell types. STIM-upregulated genes were selected using adjusted P <= 0.05 and log2 fold-change > 0.25; the top 150 all-PBMC genes were used for the global ORBIT analysis, and cell-type-specific lists were capped for downstream recurrence analysis. ORBIT output tables were used to summarize enriched terms, cell-type recurrence of enriched biology, and gene-level functional classes.

#### TCGA-BRCA tumor-subtype use case

To evaluate ORBIT in a cancer-relevant bulk RNA-seq setting, we analyzed public TCGA-BRCA expression and clinical data from UCSC Xena. PAM50 Basal tumors were compared with Luminal A/B tumors using gene-wise Welch tests on log2 expression values followed by Benjamini-Hochberg correction. The top locally annotatable basal-up and luminal-up genes were passed through the ORBIT/PANTHER-backed local annotation layer for functional-class annotation and hypergeometric enrichment against the same local annotation universe used for the figure-generation workflow.

## DECLARATIONS

### ETHICS APPROVAL AND CONSENT TO PARTICIPATE

Not applicable

### CONSENT FOR PUBLICATION

All authors have read and approved the final version of this manuscript.

### AUTHORS’ CONTRIBUTIONS

B.L.K. conceptualized the study, designed and supported the work, performed data analysis, generated the computational code, and wrote the manuscript.

## ACKNOWLEDGEMENTS

We also thank the Wayne State University High Performance Computing Grid (https://www.grid.wayne.edu/) for access to the computational resources.

## RESULTS

### ORBIT generates annotation-aware enrichment summaries from gene sets

We developed ORBIT as an annotation-aware gene-set interpretation workflow that converts input gene sets into ranked enrichment terms, semantic neighborhoods, representative functional summaries, and reviewable output tables and figures. ORBIT first maps query genes to a human annotation background, integrates protein identifiers, Gene Ontology, UniProt, Human Protein Atlas, and related evidence, and then evaluates enrichment using an empirical background-matched framework. The enriched terms are subsequently reranked using semantic coherence, gene-term alignment, and query-level functional consensus, followed by redundancy-aware representative-term selection.

This workflow was designed to address a practical limitation of standard enrichment analysis: long ranked tables often contain overlapping terms that require manual consolidation before biological interpretation. ORBIT therefore reports not only enriched terms, but also functional-class annotations, supporting-gene evidence, semantic clusters, and representative summaries that preserve an auditable path from input genes to biological interpretation (**Figure 1A**).

**Figure 1.**
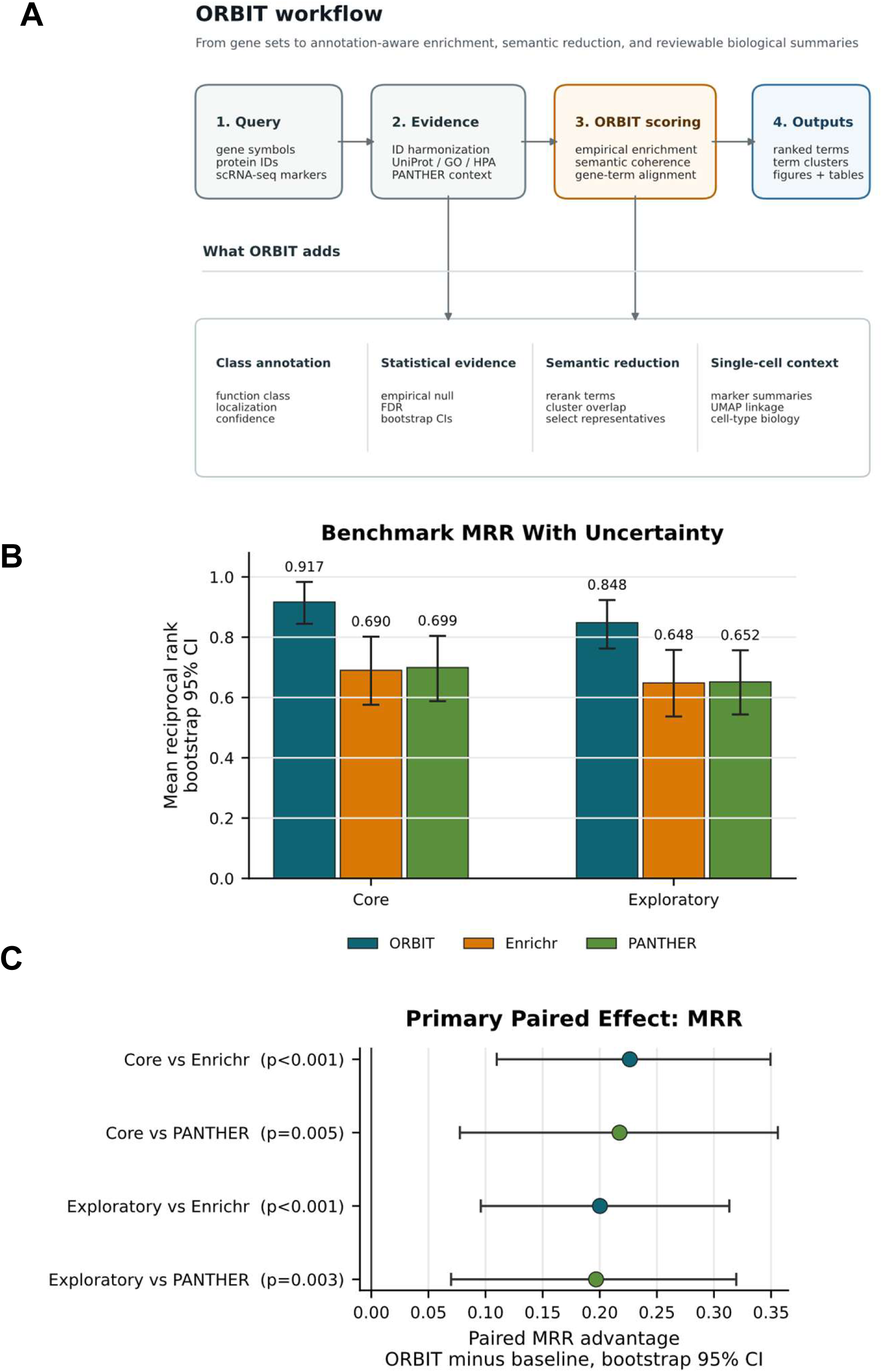
ORBIT workflow and primary benchmark performance. (A) Overview of the ORBIT workflow, from query gene sets or single-cell marker lists through evidence integration, empirical enrichment, semantic scoring, representative-term selection, and report generation. The interpretability layer summarizes ORBIT outputs used in this study: class annotation, statistical support, semantic reduction, and single-cell context. (B) Mean reciprocal rank (MRR) for ORBIT semantic, Enrichr Gene Ontology molecular-function, and PANTHER Gene Ontology molecular-function baselines across the core and exploratory benchmark scopes. Error bars show bootstrap 95% confidence intervals across benchmark gene sets. (C) Paired MRR advantage of ORBIT semantic over the two external molecular-function baselines. Points show mean paired deltas, horizontal bars show bootstrap 95% confidence intervals, and p-values are from paired sign-flip permutation tests.

### ORBIT improves functional-class recovery

We evaluated ORBIT on a curated benchmark designed to measure recovery of expected simplified functional classes from human gene sets. The benchmark tested whether annotation-aware enrichment with semantic reranking improved rank-based recovery relative to external molecular-function-oriented baselines. This evaluation emphasized first-rank and top-k recovery because practical gene-set interpretation often depends on whether the dominant biological identity of a query is captured among the first few reported terms, rather than buried lower in a long enrichment table.

Across the 45-set core benchmark, ORBIT semantic achieved the strongest expected functional-class recovery, with an MRR of 0.916, top-1 recovery of 0.889, top-5 recovery of 0.978, and top-10 recovery of 1.000. The best-performing Enrichr and PANTHER molecular-function baselines had lower MRR values of 0.690 and 0.699, respectively, and both had top-1 recovery of 0.578. Bootstrap confidence intervals and paired gene-set-level tests supported this aggregate result, with MRR used as the primary paired-effect metric (Figure 1B-C). On the core benchmark, ORBIT improved MRR over Enrichr by 0.226 (95% CI 0.110 to 0.349; paired permutation p=0.00046) and over PANTHER by 0.217 (95% CI 0.078 to 0.356; p=0.00456). Tier-stratified MRR is shown in **Supplementary Figure S1A**, and expanded paired-effect analyses across additional recovery metrics are shown in **Supplementary Figure S1B**. These paired estimates indicate that the performance difference was present at the matched gene-set level rather than arising only from aggregate summary statistics.

Comparison across ORBIT configurations showed that the semantic layer contributed most of the recovery gain. ORBIT classical ORA, the standard over-representation implementation, achieved an MRR of 0.754 and top-1 recovery of 0.644, while ORBIT empirical, which uses the background-matched empirical enrichment backbone, achieved an MRR of 0.694 and top-1 recovery of 0.533. Adding semantic reranking and representative-term selection increased performance to the full ORBIT semantic result (**Supplementary Figure S2A**). Exploratory best-rank counts and tier-stratified core benchmark MRR are shown in **Supplementary Figure S2B**-**C**.

When the benchmark was extended to the 52-set exploratory tier, ORBIT preserved the same performance advantage. ORBIT semantic achieved an MRR of 0.851 and top-1 recovery of 0.788, compared with MRR values of 0.648 and 0.652 and top-1 recovery of 0.538 for the Enrichr and PANTHER molecular-function baselines. Paired analysis again favored ORBIT, with MRR improvements of 0.200 over Enrichr (95% CI 0.096 to 0.313; p=0.00062) and 0.197 over PANTHER (95% CI 0.070 to 0.320; p=0.00344). Although smaller benchmark tiers had wider confidence intervals, particularly the seven-set secondary-reference tier, ORBIT had the highest point-estimate MRR in each tier (**Supplementary Figure S1A**).

### External comparator performance depends on enrichment library

External comparator performance varied by enrichment library and annotation source. In the benchmarked external comparisons, the strongest Enrichr baseline retained for primary analysis was GO Molecular Function 2025, while the most informative direct PANTHER outputs were PANTHER GO Molecular Function and PANTHER Protein Class (**Supplementary Figure S2A**). These results support the use of molecular-function-oriented libraries as the primary external baselines for this simplified class-recovery benchmark and are consistent with prior work showing that enrichment performance depends strongly on library selection, pathway representation, and redundancy among tested categories^10, 11, 21, 33^.

The expanded benchmark summary showed that ORBIT semantic most frequently achieved the best recovered rank across the curated benchmark sets, indicating that the observed performance gain was not limited to a small subset of queries (**Supplementary Figure S2B**). In the core benchmark, tier-stratified MRR comparisons across the broader method set showed the same overall pattern, with ORBIT semantic achieving the highest point-estimate MRR across benchmark tiers (**Supplementary Figure S2C**).

To broaden the external comparison beyond Enrichr and PANTHER, we also evaluated g:Profiler on the same tiered benchmark gene sets. ORBIT retained the highest mean reciprocal-rank recovery compared with both g:Profiler Gene Ontology molecular-function and all-source outputs (**Supplementary Figure S3A**). Paired MRR comparisons and top-k recovery summaries for the expanded comparator set are shown in **Supplementary Figure S3B**-C.

### ORBIT resolves a coherent GPCR signaling landscape from a mixed-function query

To evaluate interpretability beyond aggregate benchmark metrics, we analyzed the human GPCR mixed-hard case study. This query contained 12 input genes, produced 168 tested terms, and yielded 52 terms passing false-discovery-rate control. ORBIT recovered a coherent functional landscape centered on GPCR activity, G protein-coupled receptor activity, transmembrane receptor signaling, neurotransmitter receptor activity, chemokine-related annotations, and related receptor-ligand terms (**Figure 2**).

**Figure 2.**
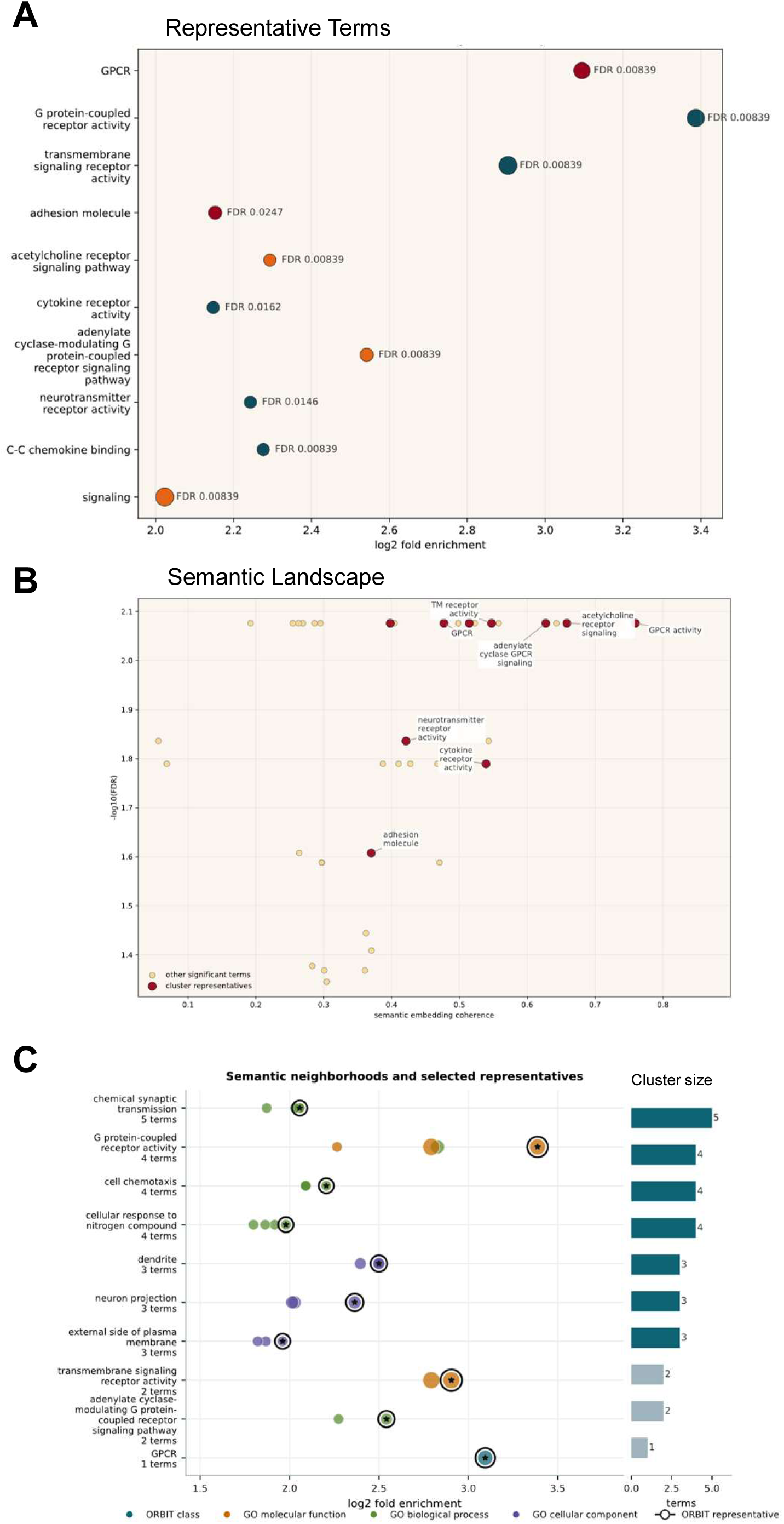
GPCR mixed-hard case study and semantic-neighborhood reduction. (A) Representative enriched terms recovered from the GPCR mixed-hard query, plotted by log2 fold enrichment. (B) Semantic landscape of significant enriched terms, showing representative terms in relation to semantic coherence and statistical significance. (C) Semantic-neighborhood reduction plot showing redundant enriched terms grouped into representative neighborhoods. Each row is a semantic neighborhood; outlined points indicate nonredundant representative terms retained by ORBIT, and right-side bars show the number of terms assigned to each neighborhood.

The representative-term plot showed that high-ranking cluster representatives captured receptor signaling and GPCR-related categories (**Figure 2A**). The semantic landscape placed these representative terms within a high-significance, high-coherence region relative to other significant terms (**Figure 2B**). The semantic-neighborhood panel then showed how related enriched terms were grouped into representative neighborhoods and which terms were retained as nonredundant ORBIT representatives (**Figure 2C**). For the same case study, supporting-gene overlap among retained representatives is shown in **Figure 3A**, and semantic-neighborhood size summaries are shown in **Figure 3B**.

**Figure 3.**
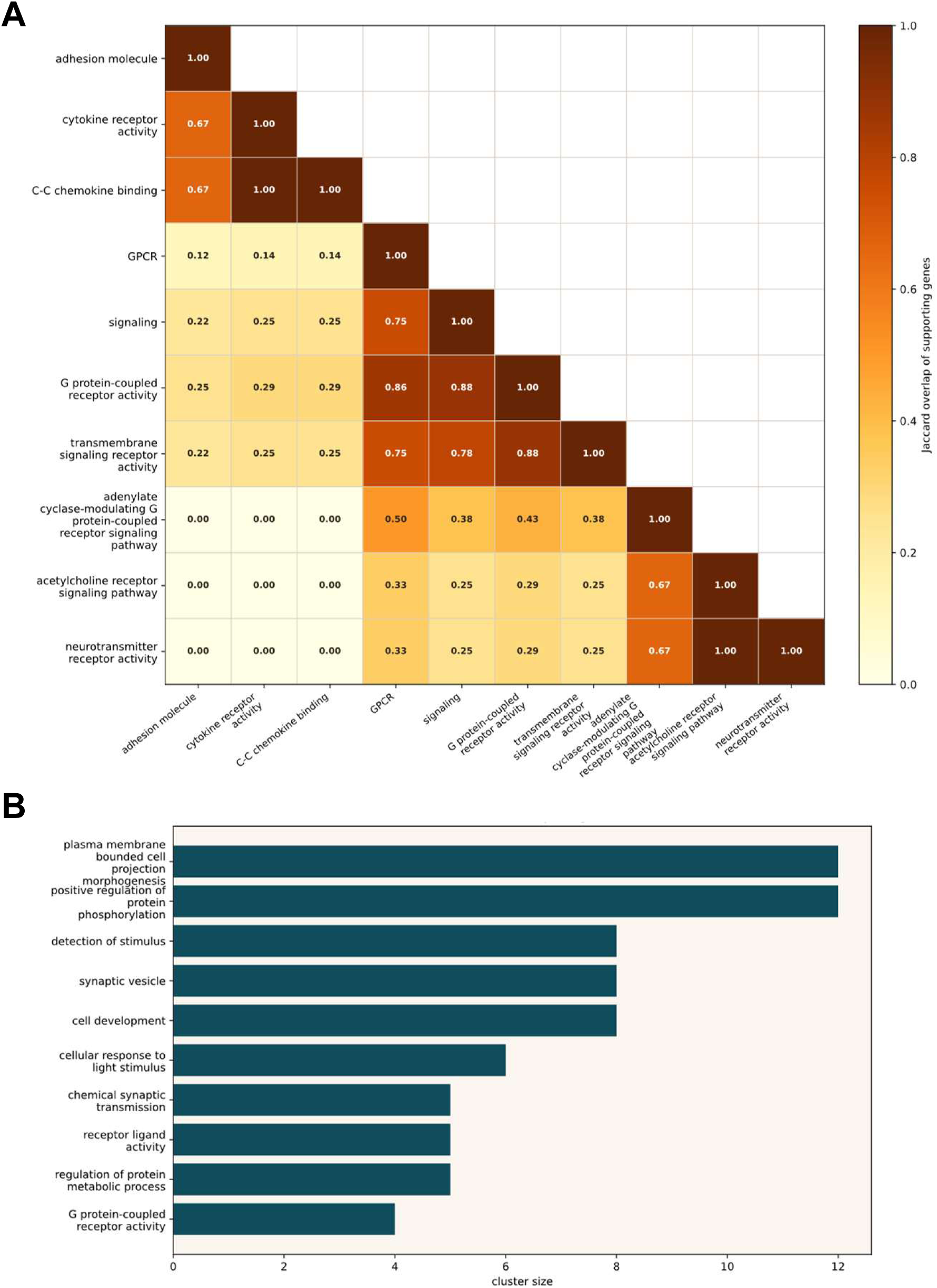
GPCR representative-term support and semantic-cluster structure. (A) Jaccard (A) Jaccard overlap among supporting query-gene subsets for hierarchically ordered GPCR case-study representative terms. The heatmap shows related supporting-gene structure among retained representatives without complete duplication across all terms. (B) Largest semantic clusters in the GPCR mixed-hard case study, summarizing the number of significant terms grouped under major representative neighborhoods.

To quantify redundancy reduction, we summarized the GPCR mixed-hard output before and after ORBIT semantic representative selection. The 52 significant enriched terms were reduced to 25 semantic representatives (**Supplementary Figure S4A**), and the largest semantic neighborhoods showed how related enriched terms were collapsed into compact representative groups (**Supplementary Figure S4B**). These results show that ORBIT condensed overlapping enrichment results while preserving interpretable neighborhood labels.

### Semantic clustering reduces redundancy across curated functional-class examples

We next evaluated ORBIT across four curated functional-class examples spanning GPCR, transcription factor, kinase, and secreted factor biology. ORBIT recovered the expected top semantic term in each case. The GPCR example returned GPCR as the top semantic term, with 43 significant terms organized into 21 semantic clusters. The transcription-factor example returned transcription factor as the top semantic term, with 19 significant terms and 13 semantic clusters. The kinase example returned kinase as the top semantic term, with 45 significant terms and 17 semantic clusters. The secreted-factor example returned secreted factor as the top semantic term, with 71 significant terms and 25 semantic clusters.

These examples illustrate how semantic clustering separates functional recovery from redundancy in enrichment outputs. Raw significant-term counts can overstate apparent biological breadth when multiple enriched terms are supported by identical or highly overlapping input-gene subsets. In the transcription-factor example, several high-overlap terms were supported by identical gene sets or by a single dominant supporting gene, producing repeated Jaccard overlap values of 1.0 in the overlap heatmap. Semantic-cluster summaries therefore provided a more compact description of the recovered functional landscape than significant-term counts alone.

### Semantic reranking improves expected-class prioritization

Together, the benchmark and case-study results indicate that ORBIT’s main gain arises from improved prioritization within enriched functional neighborhoods. The empirical enrichment backbone often placed the expected functional class in or near the top-ranked neighborhood, while semantic reranking more consistently promoted the interpretable class-level term to the top of the ranked output. This distinction is important for practical gene-set interpretation, where users often review only the first few reported terms. ORBIT’s improvement is therefore aligned with prior arguments that enrichment workflows should be evaluated not only by significance testing, but also by their ability to support coherent biological interpretation and redundancy reduction in the final output ^32, 34, 36^.

### ORBIT summarizes single-cell PBMC marker biology

We next evaluated whether ORBIT could summarize marker genes from annotated single-cell clusters. In the SeuratData PBMC3K analysis, Seurat-style UMAP annotations provided the cell-type context for nine PBMC populations (**Figure 4A**). ORBIT localization summaries showed how marker genes from each cell type mapped to broad subcellular annotation classes (**Figure 4B**), while function-class enrichment highlighted expected cell-type biology across the marker sets (**Figure 4C**). NK and CD8 T-cell marker sets were enriched for cytotoxicity-related annotations, B-cell and dendritic-cell markers recovered antigen-presentation and receptor-associated terms, monocyte markers recovered innate-immunity and calcium-binding or enzyme-associated terms, and platelet markers recovered extracellular-matrix, junction-protein, and chemokine-associated annotations. These results show that ORBIT can convert cluster marker genes into compact biological annotations and cell-type-by-function graphics that can be reviewed alongside UMAP-based cell labels. The supporting PBMC function-class heatmap and marker-gene effect-size plot are provided in **Supplementary Figure S5** and **Supplementary Figure S6**.

**Figure 4.**
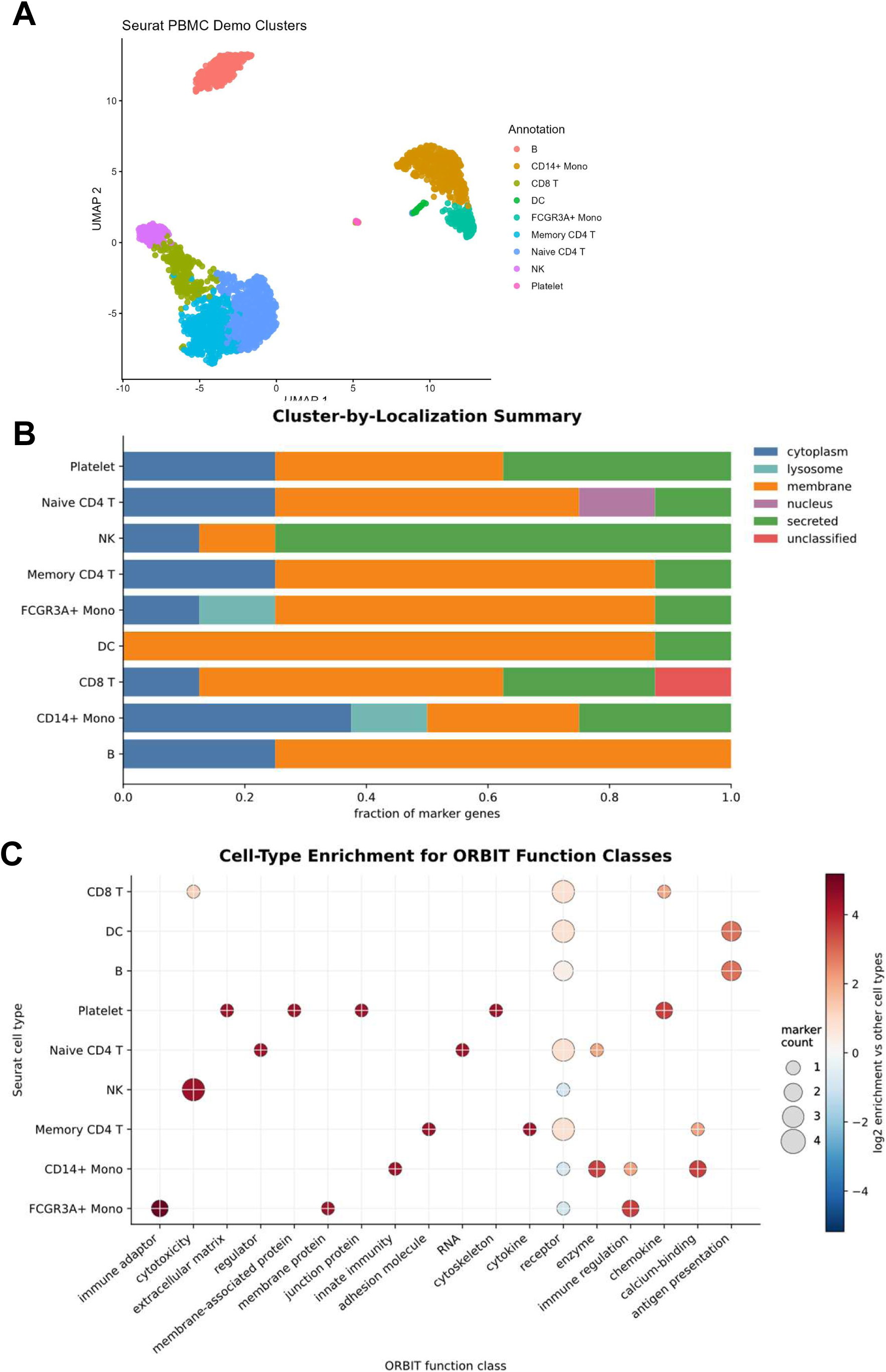
SeuratData PBMC3K single-cell marker interpretation with ORBIT. (A) UMAP of annotated PBMC3K cell populations used to define marker-gene inputs. (B) Cluster-by-localization summary showing the fraction of marker genes assigned to major localization categories for each annotated cell type. (C) Cell-type enrichment dot plot for ORBIT function classes. Dot color indicates log2 enrichment relative to other cell types, and dot size indicates the number of marker genes supporting each cell-type/function-class combination.

### ORBIT recovers perturbational IFNB-response biology from PBMC expression signatures

To evaluate ORBIT in a perturbational expression-analysis setting, we applied ORBIT to STIM-upregulated genes from the SeuratData IFNB PBMC dataset. Differential-expression results identified the DE-derived ORBIT input genes, including canonical interferon-response genes and chemokines (**Figure 5A**). ORBIT recovered antiviral and cytokine-associated biology, including defense response to virus, response to virus, double-stranded RNA binding, innate immune response, adenylyltransferase activity, RNA binding, and secreted-factor annotations (**Figure 5B**). These enriched terms recurred across major PBMC populations, indicating that ORBIT summarized a shared interferon-response program rather than a cell-type-restricted signal (**Figure 5C**). Gene-level functional-class summaries showed that the IFNB-response input genes were enriched for enzyme, secreted-factor, transcription-factor, RNA-binding, and transmembrane-protein annotations (**Figure 5D**).

**Figure 5.**
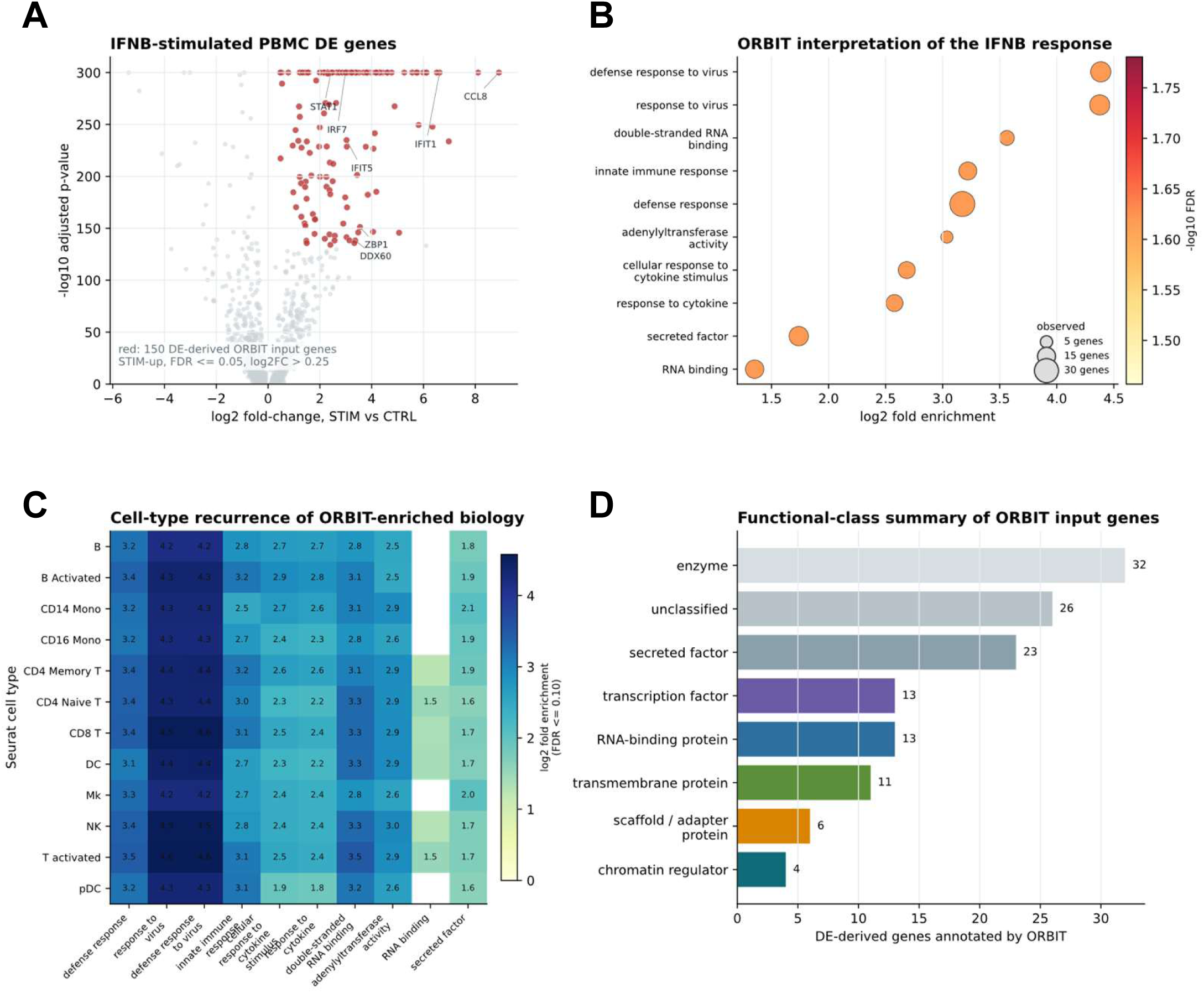
ORBIT interpretation of IFNB-stimulated PBMC expression response. (A) Volcano plot of differential expression between IFNB-stimulated and control PBMCs from the SeuratData IFNB dataset. Red points indicate the 150 STIM-upregulated genes selected as ORBIT inputs. (B) ORBIT-enriched terms from the STIM-upregulated input genes, including antiviral-response, cytokine-response, double-stranded RNA-binding, adenylyltransferase-activity, RNA-binding, and secreted-factor annotations. Dot size indicates observed input-gene count, and color indicates enrichment strength. (C) Recurrence of selected ORBIT-enriched terms across Seurat-annotated PBMC populations. (D) ORBIT gene-level functional-class summary for the DE-derived input genes.

### ORBIT identifies tumor-subtype-associated biology in TCGA-BRCA

Finally, we evaluated ORBIT in a disease-relevant bulk RNA-seq setting by analyzing TCGA-BRCA basal-like tumors relative to Luminal A/B tumors. The differential-expression signature separated basal-up and luminal-up genes that were suitable for downstream ORBIT interpretation (**Figure 6A**). ORBIT prioritized a basal-up program dominated by mitotic cell-cycle, chromosome, and centromere-associated biology, consistent with the proliferative character of basal-like breast cancer, whereas luminal-up genes were enriched for transporter and receptor-associated annotations that aligned with a more differentiated luminal epithelial state (**Figure 6B**). Gene-level functional-class summaries further showed that the two subtype signatures differed not only in individual marker genes but also in the kinds of annotated proteins contributing to each signature (**Figure 6C**). The marker-direction heatmap provided an internal biological check on the contrast: basal-associated genes including KRT5, KRT14, KRT17, EGFR, and FOXC1 were higher in basal tumors, whereas ESR1, PGR, FOXA1, GATA3, and XBP1 were higher in luminal tumors (**Figure 6D**). Thus, ORBIT converted a tumor subtype expression contrast into an interpretable hierarchy linking subtype markers, enriched biological themes, and functional protein classes.

**Figure 6.**
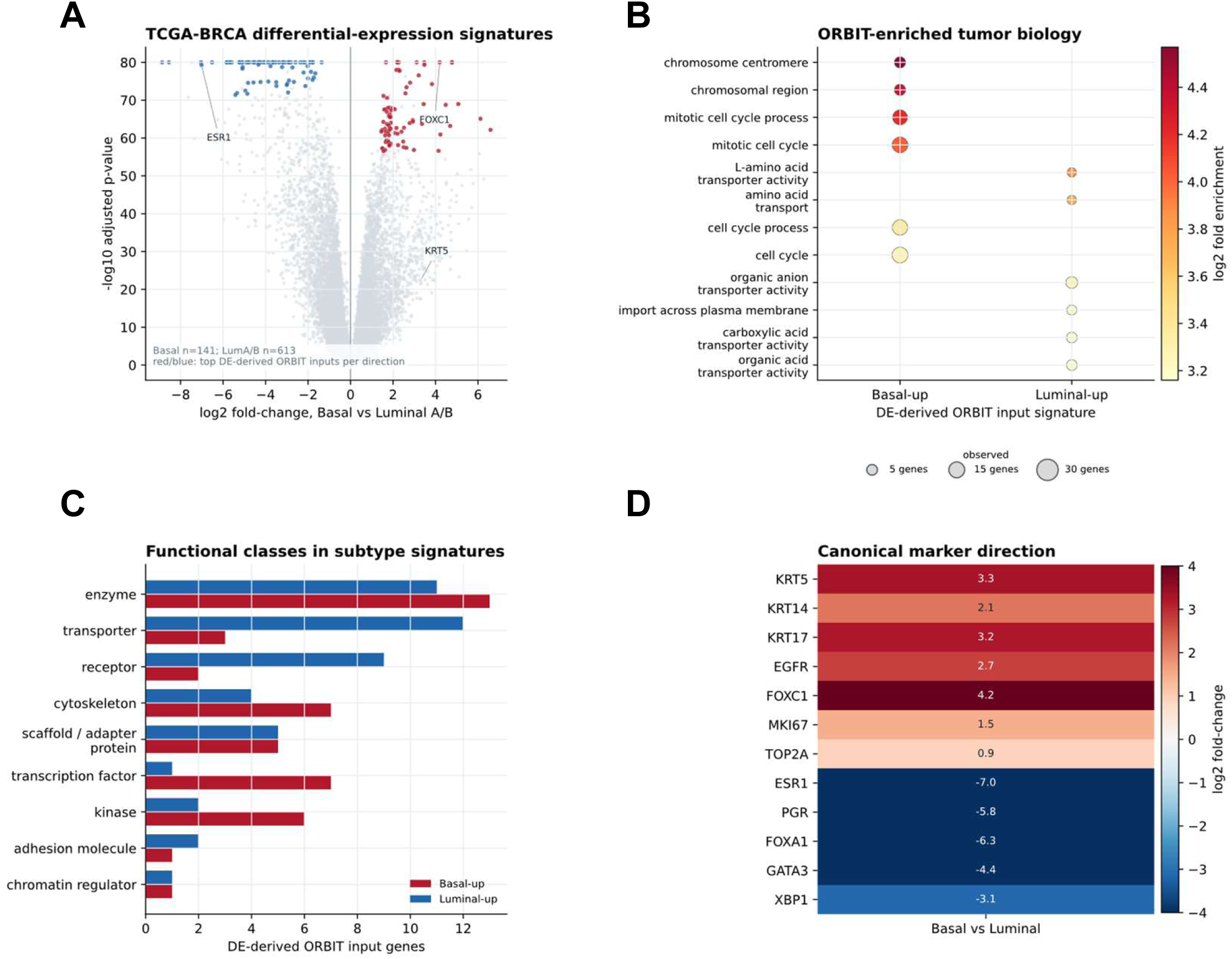
TCGA-BRCA basal-like versus luminal tumor-subtype use case. (A) Volcano plot comparing TCGA-BRCA PAM50 basal-like tumors with Luminal A/B tumors using public UCSC Xena expression and clinical annotations. Colored points indicate the DE-derived basal-up and luminal-up input genes passed to ORBIT. (B) ORBIT-enriched tumor-subtype-associated biology for basal-up and luminal-up signatures. Dot size indicates observed input-gene count, and color indicates log2 fold enrichment. (C) ORBIT simplified functional classes among subtype input genes with assigned class labels; genes without a confident simplified class assignment are omitted from this plotted summary. (D) Marker-direction heatmap showing canonical basal and luminal marker expression patterns in the analyzed contrast.

## DISCUSSION

These results support ORBIT as an annotation-aware enrichment workflow that improves recovery of interpretable functional classes from curated gene sets. Across both the 45-set core benchmark and the 52-set exploratory benchmark, ORBIT semantic outperformed the strongest tested Enrichr and direct PANTHER molecular-function baselines. The improvement was especially clear for first-rank recovery, a practical target for users who need the top returned terms to capture the dominant functional identity of a query gene set. This emphasis on rank position and interpretability complements prior discussions of gene-set analysis, which have highlighted the importance of ranking stability, category relevance, and biological reviewability in addition to nominal significance alone ^20, 21^.

This performance pattern is consistent with the design logic of ORBIT. Classical over-representation statistics can identify a relevant functional neighborhood, but statistical enrichment alone does not always determine which term within that neighborhood provides the most interpretable class-level summary. ORBIT addresses this by combining an empirical enrichment backbone with semantic coherence features and query-level consensus signals. The results suggest that semantic reranking improves placement of the expected functional class by prioritizing terms that are both statistically supported and semantically aligned with the query. This two-stage structure also aligns with prior work showing that overlap-aware or model-based gene-set interpretation can provide more concise summaries than term-by-term testing alone ^24, 33, 37^.

The benchmark also highlights the importance of comparator and library selection. Enrichr, PANTHER, and g:Profiler are valuable external resources, but their performance on this benchmark depended on the selected ontology branch, library, and output source. Molecular-function-oriented baselines were the most relevant comparators for simplified functional-class recovery, whereas broader or less directly matched libraries were less informative for this specific task. This reinforces best-practice guidance that interpretation quality depends not only on the enrichment method, but also on library relevance, term redundancy, and how outputs are summarized for downstream use ^9–11, 16–22^.

The GPCR mixed-hard case study provided a complementary interpretability result. Rather than returning an unstructured list of receptor-signaling-related terms, ORBIT organized the output into representative semantic neighborhoods that highlighted GPCR activity, receptor signaling, neurotransmitter receptor activity, chemokine-related signaling, and related functional themes. This representation supports direct review of high-ranking representatives, semantic coherence, supporting genes, and neighborhood-level reduction rather than requiring users to infer structure from dozens of partially redundant enrichment hits (**Figure 2**; **Figure 3**). This emphasis on summarization rather than raw list length is consistent with previous observations that GO-based enrichment frequently returns correlated terms and benefits from grouping, clustering, or ontology-aware abstraction layers^23, 33, 37^.

The curated functional-class examples broadened the interpretability result across distinct annotation structures. ORBIT recovered the expected top semantic term for GPCR, transcription-factor, kinase, and secreted-factor inputs. The transcription-factor case was especially informative because several apparently distinct ontology terms were supported by the same input-gene subset or by a single dominant supporting gene. This overlap structure illustrates why representative-term clustering is useful: raw significant-term counts can overstate apparent biological breadth, whereas semantic cluster counts and representative terms provide a more compact summary of the recovered functional landscape. Similar concerns about inflated result lists and the need for higher-level summarization recur throughout the enrichment literature, including ontology review papers, semantic-similarity tool papers, and practical pathway-analysis guidelines ^23, 24, 34^.

A residual limitation was observed in enzyme-class benchmark sets. In these cases, ORBIT often prioritized specific catalytic subtypes over broader generic enzyme labels. This behavior can be biologically informative because specific catalytic terms may provide more mechanistic detail, but it can reduce expected-label recovery when the benchmark target is defined at a coarser class level. Future development should therefore add explicit control over functional granularity, allowing users to tune whether ORBIT favors broad class-level summaries or more specific mechanistic terms.

Several considerations should guide interpretation of these results. The benchmark was designed to evaluate recovery of simplified functional classes from curated human gene sets, and the strongest claims therefore apply to this class-recovery setting. The primary comparisons focused on molecular-function-oriented Enrichr and PANTHER baselines, with g:Profiler included as an additional external comparator. Applied PBMC3K, IFNB, and TCGA-BRCA analyses should be interpreted as use-case demonstrations rather than independent statistical benchmarks. Overall, the strongest support for ORBIT comes from the matched benchmark with bootstrap confidence intervals, paired gene-set-level tests, and transparent supplemental tables, while the applied transcriptomic examples illustrate how the workflow can translate marker and differential-expression signatures into structured biological interpretation.

## CONCLUSIONS

ORBIT was developed to improve gene-set interpretation by combining empirical enrichment, semantic reranking, and representative-term clustering within an annotation-aware workflow. On the curated tiered benchmark evaluated here, ORBIT semantic achieved higher recovery of expected functional-class labels than the tested Enrichr and PANTHER molecular-function baselines across both core and exploratory benchmark scopes, with supplemental g:Profiler comparisons showing the same overall pattern.

Across the GPCR mixed-hard case study, additional curated functional-class examples, and applied PBMC3K, IFNB, and TCGA-BRCA analyses, ORBIT produced compact representative-term summaries with traceable supporting genes and functional classes. Together, these findings support ORBIT as a practical workflow for improving rank-based functional-class recovery and reducing redundancy in enrichment outputs.

## Availability and implementation

ORBIT is implemented as the Python package orbit, version 0.1.0, with the command-line entry point orbit. The package is released under the MIT license and supports Python >=3.11. The version used for this manuscript is available as the ORBIT v0.1.0 GitHub release at https://github.com/KidderLab/ORBIT/releases/tag/v0.1.0 and is archived at Zenodo with DOI 10.5281/zenodo.20849753.

Installation from a source checkout can be performed with python -m pip install -e.. A minimal enrichment run is orbit enrich --input input.csv --species human --outdir results/enrichment --permutations 1000, where input.csv contains a column named identifier or a first column of gene/protein identifiers. The PBMC3K single-cell demonstration can be reproduced with orbit pbmc-demo --outdir results/pbmc_demo.

The manuscript analysis bundle includes benchmark input manifests, ORBIT result tables, Enrichr, PANTHER, and g:Profiler comparator outputs, PBMC3K marker outputs, IFNB and TCGA-BRCA use-case outputs, figure-generation scripts, and supplemental tables. Outputs are provided in standard formats, including CSV, JSON, HTML, PNG/SVG, and Excel. ORBIT can run annotation-aware enrichment locally from offline background resources, while optional live annotation and external comparator workflows depend on external APIs or web services. Unmapped genes are retained in query annotation tables with mapping-status fields so mapping failures can be audited rather than silently removed.

