## Supplementary Information for "ORBIT: Annotation-Aware Empirical Enrichment and Semantic Reranking for Interpretable Functional-Class Recovery"

Benjamin L. Kidder[1-2*]

[1]Department of Oncology, Wayne State University School of Medicine, Detroit, MI, USA

[2]Karmanos Cancer Institute, Wayne State University School of Medicine, Detroit, MI, USA

Running title: ORBIT semantic gene-set interpretation

*Correspondence:

Benjamin L. Kidder

### Supplemental Figures

**Figure S1. Benchmark robustness analyses.** (A) Mean reciprocal rank stratified by benchmark tier. Error bars show bootstrap 95% confidence intervals, and within-bar labels indicate the number of benchmark gene sets in each tier. (B) Expanded paired ORBIT advantage analysis across MRR and top-k recovery metrics. Points show mean paired deltas relative to the indicated external baseline, horizontal bars show bootstrap 95% confidence intervals, and p-values are from paired sign-flip permutation tests.

**Figure S2. Benchmark summary across ORBIT modes and external baselines**. (A) Core benchmark performance across MRR and top-k recovery metrics for ORBIT semantic, ORBIT classical ORA, ORBIT empirical, Enrichr Gene Ontology molecular-function, PANTHER Gene Ontology molecular-function, and PANTHER protein-class baselines. (B) Exploratory benchmark best-rank counts across methods. (C) Core benchmark MRR by benchmark tier across methods.

**Figure S3. Expanded external enrichment comparator analysis.** (A) Mean reciprocal-rank recovery for ORBIT, Enrichr, PANTHER, and g:Profiler comparator outputs on the core and exploratory benchmark sets. (B) Paired core-benchmark MRR advantage for ORBIT relative to each external comparator, with bootstrap 95% confidence intervals. (C) Core benchmark top-k recovery rates for the same comparator set.

**Figure S4. Semantic-reduction audit for ORBIT redundancy control.** (A) Number of significant enriched terms before and after semantic representative selection in the GPCR mixed-hard case study. (B) Largest semantic neighborhoods collapsed by ORBIT into representative terms.

**Figure S5. PBMC clustered cell-type by function-class heatmap.** Marker-gene function-class fractions are shown for each Seurat-annotated PBMC cell type. Values indicate the fraction of marker genes assigned to each ORBIT function class, highlighting broad cell-type-specific annotation patterns that complement the enrichment dot plot in Figure 4.

**Figure S6. PBMC marker-gene effect-size and prevalence plot.** Top marker genes for each annotated PBMC cell type are shown by rank within cell type. Color indicates average log2 fold-change, and point size indicates marker prevalence or supporting-cell fraction, depending on the plotted variable. Marker labels identify genes supporting the cell-type annotation patterns summarized in Figure 4 and Supplementary Figure S5.

A

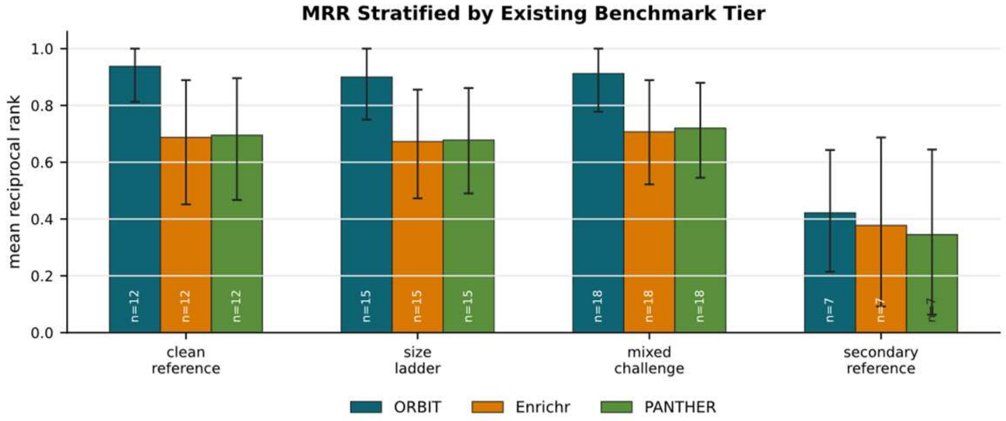
MRR Stratified by Existing Benchmark Tier
mean reciprocal rank
1.0
0.8
0.6
0.4
0.2
0.0
n=12
n=15
n=18
n=7
clean reference
size ladder
mixed challenge
secondary reference
ORBIT
Enrichr
PANTHER

B

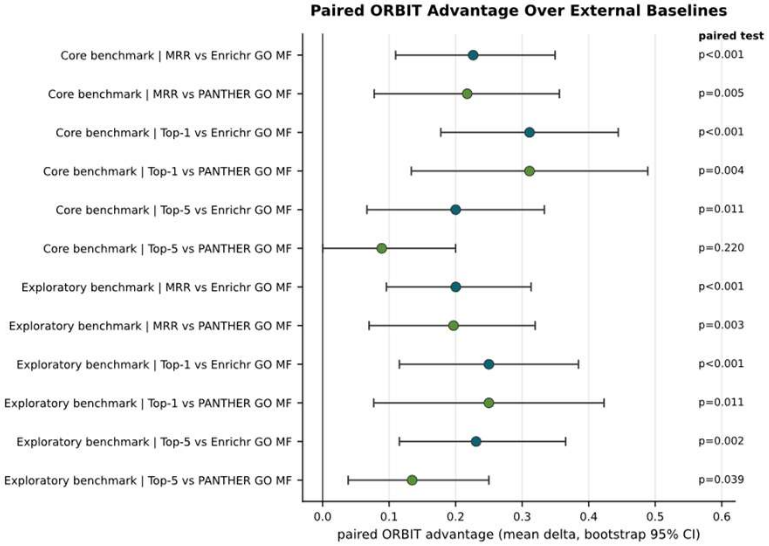
Paired ORBIT Advantage Over External Baselines
paired test
Core benchmark | MRR vs Enrichr GO MF
p<0.001
Core benchmark | MRR vs PANTHER GO MF
p=0.005
Core benchmark | Top-1 vs Enrichr GO MF
p<0.001
Core benchmark | Top-1 vs PANTHER GO MF
p=0.004
Core benchmark | Top-5 vs Enrichr GO MF
p=0.011
Core benchmark | Top-5 vs PANTHER GO MF
p=0.220
Exploratory benchmark | MRR vs Enrichr GO MF
p<0.001
Exploratory benchmark | MRR vs PANTHER GO MF
p=0.003
Exploratory benchmark | Top-1 vs Enrichr GO MF
p<0.001
Exploratory benchmark | Top-1 vs PANTHER GO MF
p=0.011
Exploratory benchmark | Top-5 vs Enrichr GO MF
p=0.002
Exploratory benchmark | Top-5 vs PANTHER GO MF
p=0.039
0.0 0.1 0.2 0.3 0.4 0.5 0.6
paired ORBIT advantage (mean delta, bootstrap 95% CI)

**Figure S1**

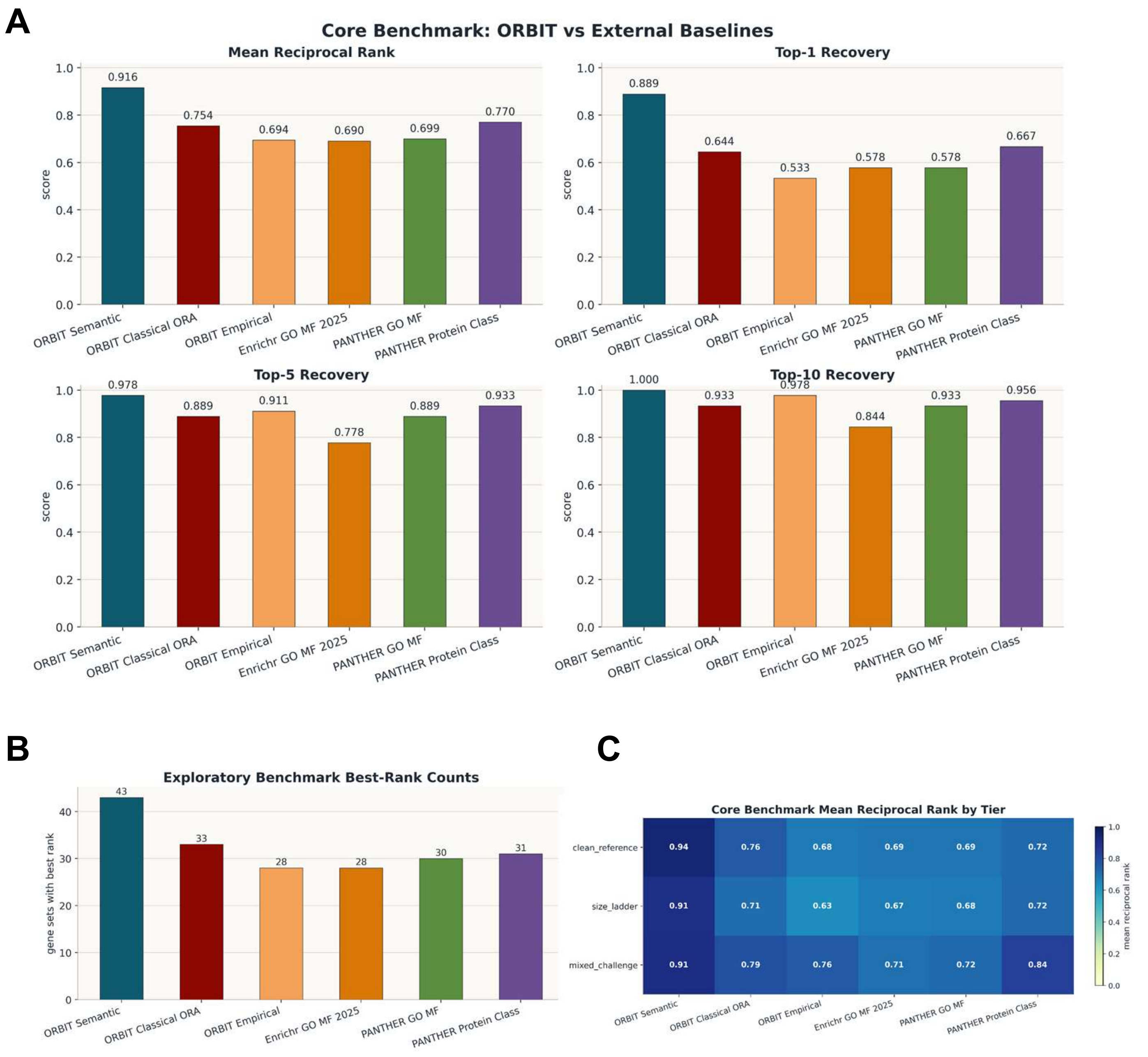
A
Core Benchmark: ORBIT vs External Baselines
Mean Reciprocal Rank
Top-1 Recovery
Top-5 Recovery
Top-10 Recovery
score
ORBIT Semantic
ORBIT Classical ORA
ORBIT Empirical
Enrichr GO MF 2025
PANTHER GO MF
PANTHER Protein Class
0.916
0.754
0.694
0.690
0.699
0.770
0.889
0.644
0.533
0.578
0.578
0.667
0.978
0.889
0.911
0.778
0.889
0.933
1.000
0.933
0.978
0.844
0.933
0.956
B
Exploratory Benchmark Best-Rank Counts
gene sets with best rank
43
33
28
28
30
31
C
Core Benchmark Mean Reciprocal Rank by Tier
clean_reference
size_ladder
mixed_challenge
0.94 0.76 0.68 0.69 0.69 0.72
0.91 0.71 0.63 0.67 0.68 0.72
0.91 0.79 0.76 0.71 0.72 0.84
mean reciprocal rank

Figure S2

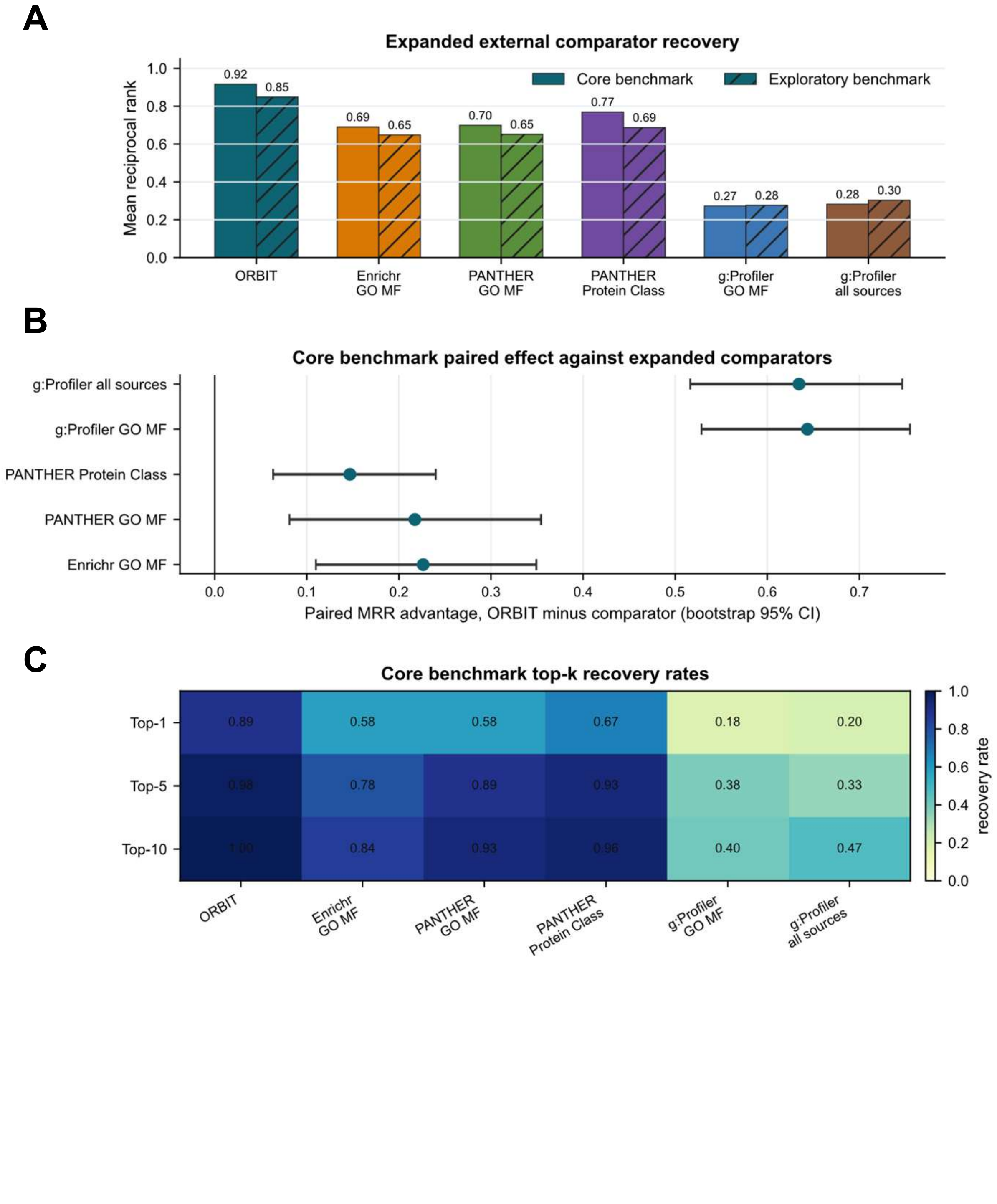
A
Expanded external comparator recovery
Mean reciprocal rank
Core benchmark
Exploratory benchmark
0.92
0.85
0.69
0.65
0.70
0.65
0.77
0.69
0.27
0.28
0.28
0.30
ORBIT
Enrichr GO MF
PANTHER GO MF
PANTHER Protein Class
g:Profiler GO MF
g:Profiler all sources
B
Core benchmark paired effect against expanded comparators
g:Profiler all sources
g:Profiler GO MF
PANTHER Protein Class
PANTHER GO MF
Enrichr GO MF
Paired MRR advantage, ORBIT minus comparator (bootstrap 95% CI)
C
Core benchmark top-k recovery rates
Top-1
Top-5
Top-10
0.89
0.58
0.58
0.67
0.18
0.20
0.98
0.78
0.89
0.93
0.38
0.33
1.00
0.84
0.93
0.96
0.40
0.47
recovery rate
ORBIT
Enrichr GO MF
PANTHER GO MF
PANTHER Protein Class
g:Profiler GO MF
g:Profiler all sources

**Figure S3**

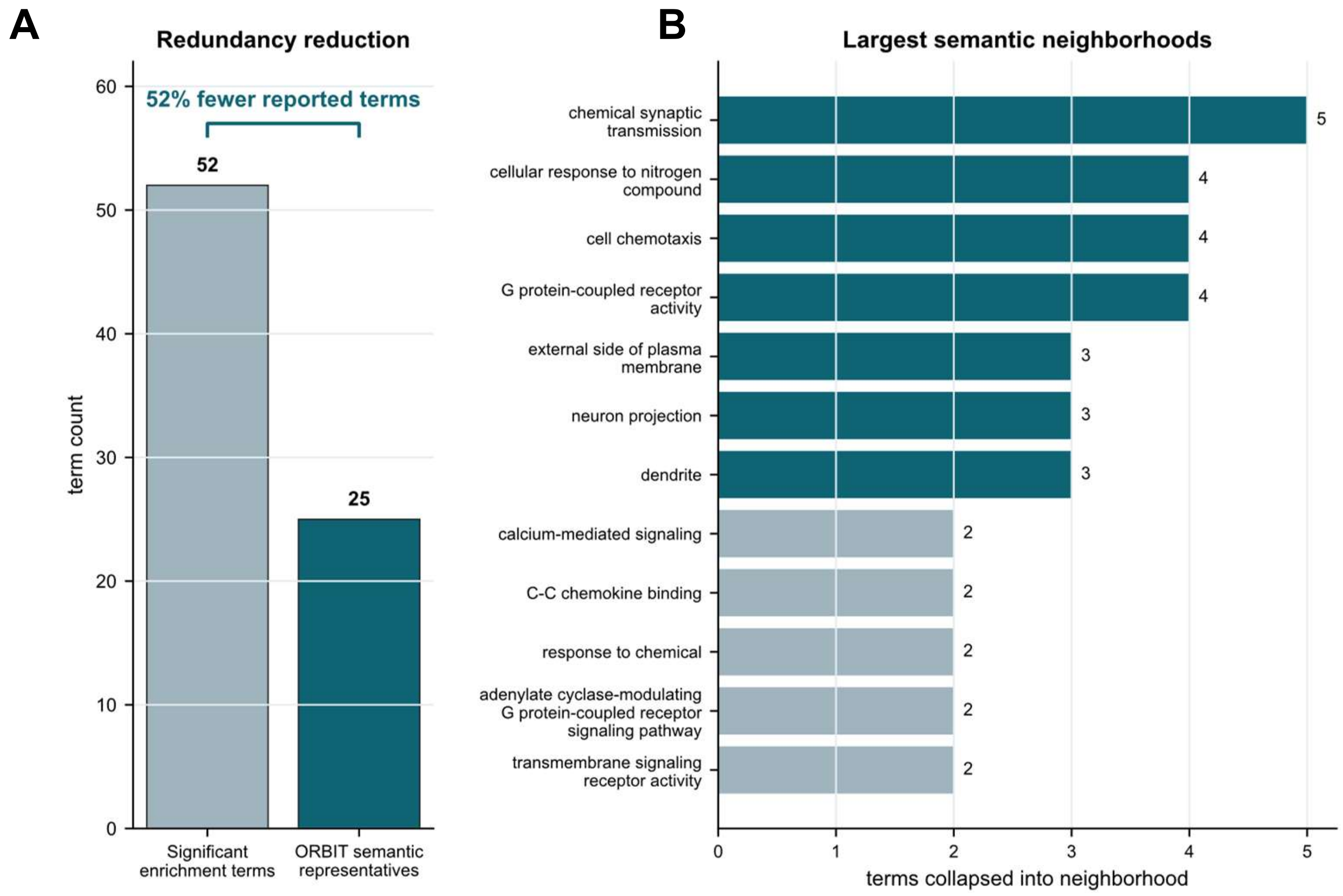
A
Redundancy reduction
52% fewer reported terms
52
25
term count
0
10
20
30
40
50
60
Significant enrichment terms
ORBIT semantic representatives
B
Largest semantic neighborhoods
chemical synaptic transmission
5
cellular response to nitrogen compound
4
cell chemotaxis
4
G protein-coupled receptor activity
4
external side of plasma membrane
3
neuron projection
3
dendrite
3
calcium-mediated signaling
2
C-C chemokine binding
2
response to chemical
2
adenylate cyclase-modulating G protein-coupled receptor signaling pathway
2
transmembrane signaling receptor activity
2
0
1
2
3
4
5
terms collapsed into neighborhood

Figure S4

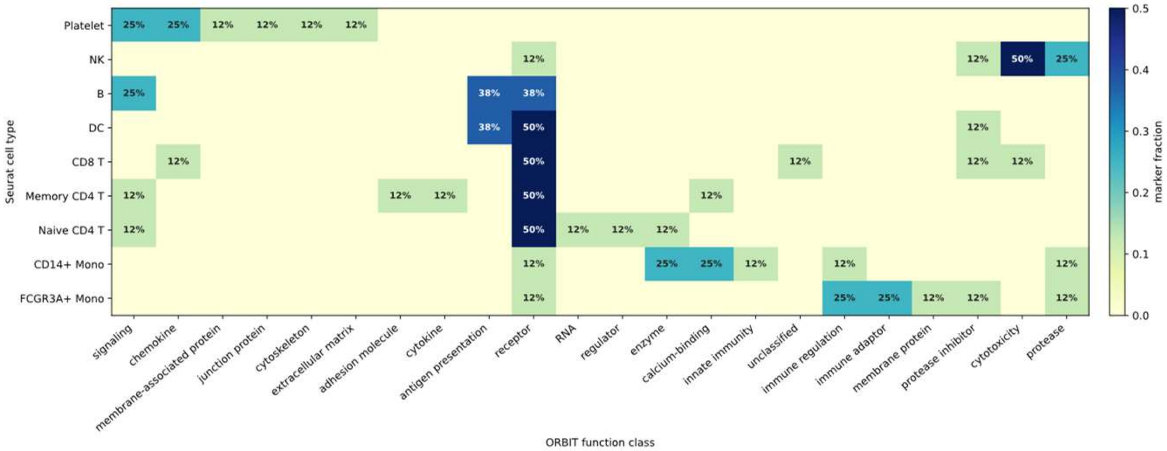

Platelet
NK
B
DC
CD8 T
Memory CD4 T
Naive CD4 T
CD14+ Mono
FCGR3A+ Mono
Seurat cell type
signaling
chemokine
membrane-associated protein
junction protein
cytoskeleton
extracellular matrix
adhesion molecule
cytokine
antigen presentation
receptor
RNA
regulator
enzyme
calcium-binding
innate immunity
unclassified
immune regulation
immune adaptor
membrane protein
protease inhibitor
cytotoxicity
protease
ORBIT function class
marker fraction


**Figure S5**

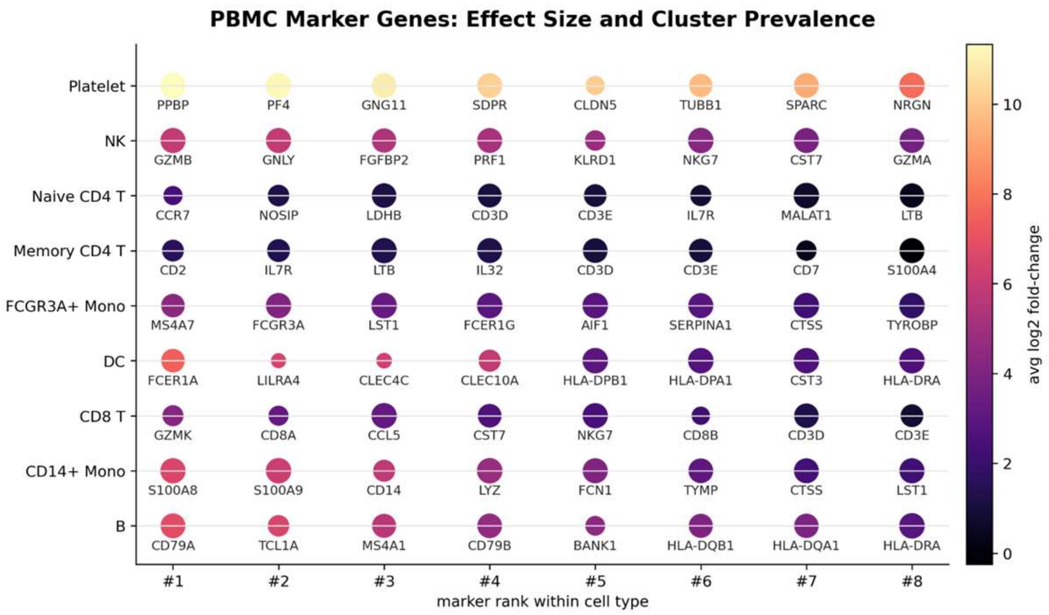
PBMC Marker Genes: Effect Size and Cluster Prevalence
Platelet
NK
Naive CD4 T
Memory CD4 T
FCGR3A+ Mono
DC
CD8 T
CD14+ Mono
B
PPBP
PF4
GNG11
SDPR
CLDN5
TUBB1
SPARC
NRGN
GZMB
GNLY
FGFBP2
PRF1
KLRD1
NKG7
CST7
GZMA
CCR7
NOSIP
LDHB
CD3D
CD3E
IL7R
MALAT1
LTB
CD2
IL7R
LTB
IL32
CD3D
CD3E
CD7
S100A4
MS4A7
FCGR3A
LST1
FCER1G
AIF1
SERPINA1
CTSS
TYROBP
FCER1A
LILRA4
CLEC4C
CLEC10A
HLA-DPB1
HLA-DPA1
CST3
HLA-DRA
GZMK
CD8A
CCL5
CST7
NKG7
CD8B
CD3D
CD3E
S100A8
S100A9
CD14
LYZ
FCN1
TYMP
CTSS
LST1
CD79A
TCL1A
MS4A1
CD79B
BANK1
HLA-DQB1
HLA-DQA1
HLA-DRA
#1
#2
#3
#4
#5
#6
#7
#8
marker rank within cell type
avg log2 fold-change
10
8
6
4
2
0

**Figure S6**
